# Role of IQGAP1 in Head and Neck Carcinogenesis

**DOI:** 10.1101/253484

**Authors:** Tao Wei, Suyong Choi, Darya Buehler, Alan C. Rapraeger, Richard A. Anderson, Paul F. Lambert

## Abstract

Head and neck squamous cell carcinoma (HNSCC) is a common cancer in humans. Phosphoinositide 3-kinase (PI3K)/AKT signaling, along with its downstream effector mTOR, is one of the most frequently altered pathways in HNSCC, and is downstream of growth factor signaling including that mediated by epidermal growth factor receptor (EGFR), which is also commonly upregulated in HNSCC. Recently, IQ motif-containing GTPase activation protein 1 (IQGAP1) has been reported to function as a scaffold for the enzymes involved in the PI3K/AKT signaling pathway. *Iqgap1* gene expression is increased in human HNSCCs, raising the hypothesis that it acts as an oncogene in this cancer. Whether IQGAP1 is necessary for HNSCC development as well as what is its underlying mechanism are both unknown. Here we report on the role of IQGAP1 in HNSCC by performing a combination of *in vitro* studies using human cancer cell lines, and *in vivo* studies using a well-validated preclinical mouse model for HNSCC that is known to depend upon EGFR signaling. Cells knocked out for IQGAP1 lost AKT signaling. Disruption of IQGAP1-scaffolded PI3K/AKT signaling using a peptide that interferes with the ability of IQGAP1 to bind to PI3K reduced HNSCC cell survival. *In vivo* studies utilizing *Iqgap1*-null (*Iqgap1*^*-/-*^) mice demonstrated that IQGAP1 is necessary for efficient PI3K signaling upon EGF-stimulation. Treatment of *Iqgap1*^*-/-*^ mice with the oral carcinogen, 4-nitroquinoline 1-oxide (4NQO), *Iqgap1*^*-/-*^ led to significantly lower multiplicities of cancer foci as well as significantly lower numbers of high grade cancers than observed in similarly treated *Iqgap1*^*+/+*^ mice. IQGAP1 protein was increased in its expression in HNSCCs arising in the *Iqgap1*^*+/+*^ mice, consistent with that seen in human HNSCCs. We also observed a significant down-regulation of PI3K signaling in 4NQO-induced HNSCCs arising in the *Iqgap1*^*-/-*^ mice, consistent with IQGAP1 contributing to carcinogenesis by promoting PI3K signaling. Our studies, therefore, support the hypothesis that IQGAP1 acts as an oncogene in head and neck carcinogenesis, and provide mechanistic insight into its role.

## INTRODUCTION

Head and neck squamous cell carcinomas (HNSCCs) arise in the squamous cells of the mouth/throat region. HNSCC the sixth most frequent cancer worldwide with a 5-year survival rate less than 50% [1]. Risk factors for HNSCC include smoking, alcohol abuse, and infection with high-risk human papillomaviruses (HPVs) [1]. Multiple studies identifying pathways that contribute to HNSCC development have observed phosphoinositide 3-kinase (PI3K) signaling to be one of the most frequently altered pathways in HNSCC patients [2-6].

PI3K is a major player in diverse physiologic processes including cell survival, migration and proliferation. In response to agonists, PI3K converts phosphatidylinositol-4,5-biphoshate (PIP_2_) to phosphatidylinositol-3,4,5-trisphosphate (PIP_3_). PIP_3_ then functions as a messenger to activate AKT, which activates to multiple downstream effectors including the mTOR1 pathway to promote cell growth and protein synthesis [7, 8]. PI3K pathway mutations occur in approximately 30% of head and neck tumors [6]. The TCGA database indicates that *PIK3CA*, the gene encoding the 110kDa catalytic subunit of PI3K, is mutated in 20% of HNSCCs patients [3]. These *PIK3CA* mutations increase PI3K activity and thereby promote HNSCC cell growth [6]. Pan-PI3K inhibitors suppress HNSCC cell growth both in tissue culture and in xenograft studies *in vivo* [8-10]. However, clinical treatment with small molecule inhibitors targeting the PI3K AKT/mTOR pathway have had limited success due to toxicity and possible drug resistance [8]. Drugs targeting other components of the PI3K/AKT/mTOR pathway in HNSCC may provide more effective treatments.

Recently, I<u>Q</u> motif-containing <u>G</u>TPase <u>a</u>ctivating <u>p</u>rotein 1(IQGAP1), has been identified as a scaffold protein that promotes PI3K/AKT/mTOR signaling [11]. IQGAP1 is a large, 190-kDa protein that contains multiple protein-interaction domains. With over 100 interacting protein partners, IQGAP1 regulates diverse cellular processes including cytoskeletal dynamics, adhesion, vesicular trafficking, cell motility/invasion, and cell proliferation [12-15]. It can also bind directly to the epidermal growth factor receptor (EGFR) and mediate receptor activation [16] as well as scaffold and facilitates the Ras-MAPK pathway [12]. IQGAP1 also binds directly to β-catenin, a key regulator of Wnt signaling, and enhances its nuclear localization and β-catenin-dependent transcription [17]. Recognized as an oncogene, IQGAP1 is overexpressed in several cancers, including those arising in the breast, lung, colorectal and head and neck region [15, 18]. Diminished tumorigenesis was observed in *Iqgap1*-knockout mice in a Ras-driven skin cancer model, consistent with its role in Ras signaling [12]. Recently, the IQGAP1 scaffold localized to cell surface receptor was reported to assemble each of the kinases, including PI3K and AKT, necessary for rapid, channeled conversion of PI to PIP3 upon receptor activation, resulting in increased AKT activation in cultured cells and *in vivo* [11]. Another study showed that, upon IQGAP1 overexpression, IQGAP1-dependent phosphorylation of ERK1/2 is impeded via AKT-mediated phosphorylation of Forkhead box protein O1 (FOXO1), a downstream effector of AKT, suggesting that there is crosstalk among IQGAP1-mediated pathways [19].

In Choi *et al*., 2016, the IQ3 motif and WW motif on IQGAP1 were identified to be the PI3K binding site, and peptides (IQ3 and WW) that contain an N-terminal, membrane-permeating motif and either the IQ3 or WW motif, were demonstrated to competitively inhibit the PI3K:IQGAP1 interaction [11]. Addition of these peptides to cells disrupted the intracellular formation of the scaffolded PI3K complex and inhibited AKT activation [11]. IQ3 and WW selectively diminished breast cancer cell survival, including cell lines harboring *PIK3CA* mutation, while not affecting normal cells [11]. This indicated that IQGAP1-mediated PI3K signaling is important for cancer cell survival.

In this study, we focus on the role of IQGAP1 in HNSCC using a well-established mouse model for HNSCC, in which a synthetic oral carcinogen, 4-nitroquinoline 1-oxide (4NQO), is used to drive tumorigenesis [20]. This carcinogen causes a spectrum of DNA damage similar to that caused by tobacco-associated carcinogens [20]. Notably, 4NQO is also known to cause HNSCC in mice at least in part by activating EGFR signaling, upstream of the PI3K/AKT/mTOR pathway, and therefore was chosen as a preclinical model that closely reflects activation of EGFR in HNSCC arising in humans [7, 20].

To test the role of IQGAP1, we used a genetic approach that makes use of a genetically modified strain of mice that is deficient for IQGAP1 [12, 21]. These *Iqgap1*- knockout *(Iqgap1*^*-/-*^) mice develop normally and have no apparent phenotypes [21]. We observed a decrease in AKT activation in *Iqgap1*^*-/-*^ mice upon EGF stimulation. Upon 4NQO treatment, we observed that loss of IQGAP1 reduced the frequency of high grade HNSCC. In the absence of IQGAP1, PI3K signaling was significantly decreased in 4NQO-induced HNSCC. In *Iqgap1*^*+/+*^ mice, IQGAP1 protein level was upregulated in HNSCC, consistent with what is observed in human HNSCCs [18]. Our studies support the hypothesis that IQGAP1 functions as an oncogene in head and neck carcinogenesis, and does so at least in part by activating the PI3K/AKT signaling pathway.

## MATERIALS AND METHODS

### Cell culture

MDA-MB-231, UPCI-SCC90 cells were purchased from ATCC and maintained in in DMEM supplemented with 10% fetal bovine serum (Gibco). UM-SCC1 cells were purchased from Millipore and maintained as described by the source.

### CRISPR-Cas9 cell line generation

To knockout *IQGAP1* gene in MDA-MB-231, CRISPR-Cas9 genome editing method was used [22]. Guide RNA sequences (5’-CCCGTCAACCTCGTCTGCGG-3’ and 5’-GGCGTGGCCCGGCCGCACTA-3’ for human *IQGAP1* gene’s 1^st^ exon) were cloned into PX459V2.4 vector. Constructs were transfected for 36 h and then transiently selected with 1 mg/ml puromycin. After 48 h incubation, puromycin was removed and single cells were seeded in 96-well tissue culture dishes. Cells were expanded and positive colonies were selected by immunoblotting with an anti-IQGAP1 antibody.

### Peptide treatment and cell counting

IQ3 and WW peptides were generated as described in [11]. HNC cells were plated in each well of 6-well dish at ∼20% confluency one day before peptide treatment. Cells were changed with fresh culture media before treating with 30μM peptides. Cells were changed with fresh media and peptide every 24 hours. After 72-hour incubation, floating cells were removed by washing with PBS and the number of viable cells was counted by staining with trypan blue.

### Mice

*Iqgap1*^*+/+*^ and *Iqgap1*^*-/-*^ mice (from Dr. David Sacks, National Institutions of Health) [21] were on a mixed genetic background (50%*FVB*/50%(*129+C57Bl/6*)). All mice were genotyped by PCR using primers from [21]. Mice were housed in the Association for Assessment of Laboratory Animal Care-approved McArdle Laboratory Animal Care Unit. All procedures were carried out in accordance with an animal protocol approved by the University of Wisconsin Institutional Animal Care and Use Committee.

### *In vivo* EGF stimulation

Treatment with epidermal growth factor (EGF) was as previously described [23]. Briefly, 5-week old *Iqgap1*^*+/+*^ and *Iqgap1*^*-/-*^ mice, were injected subcutaneously with 5ug/g (body weight) of hEGF (Prospec, CYT-217) in PBS, or PBS vehicle control. hEGF was stored as 1 mg/mL stock solution in phosphate-buffered saline (PBS) at -80˚C. Mice were sacrificed 10 minutes after injection and dorsal skin was harvested and snap frozen in liquid nitrogen.

### Protein lysate preparation and western blotting

Frozen skin was cut into small pieces on ice using a razor blade, then homogenized in 300μL RIPA buffer (25mM Tris (PH=8), 150mM NaCl, 0.1%SDS, 0.5% sodium deoxycholate, 1%Triton X-100, protease inhibitor and phosphatase inhibitor) in a 1.5mL Eppendorf tube using a plastic pestle (Axygen), and incubated on an orbital shaker at 4˚C for 20 minutes. Homogenates were centrifuged at 14,000rpm for 20 minutes at 4˚C, and resulting supernatants (protein lysates) collected. Protein concentrations were determined using the Bradford Protein Assay (Bio-Rad). Equivalent amounts of protein were resolved on precast Mini-PROTEAN TGX 7.5% gels (Bio-Rad) and transferred to PVDF membranes (Millipore). Following transfer, membranes were blocked with 5% nonfat dry milk in TBS (Tris-buffered saline with 0.1% Tween-20). Proteins were detected using anti–β-actin (1:5,000 dilution) (Sigma), anti-phospho-AKT (1:500, #9271, Cell Signaling), anti-AKT (1:1000, #9272, Cell Signaling), anti-phospho-ERK1/2 (1:1000, #4376, Cell Signaling), anti-ERK1/2 (1:1000, #4695, Cell Signaling), anti-phospho-S6 (1:1000, #4858, Cell Signaling), anti-S6 (1:1000, #2217, Cell Signaling), anti-IQGAP1 (1:1000, ab133490, Abcam), or anti-IQGAP3 (1:2000, gift from Dr. Sachiki Tsukita, Osaka University [24]) primary antibodies. Horseradish peroxidase-conjugated secondary antibodies (1:10,000) (Jackson ImmunoResearch) and chemiluminescent substrates (Clarity ECL Substrates; Bio-Rad) were used to visualize immune complexes on a Bio-Rad ChemiDoc Imaging System.

### 4-nitroquinoline-1-oxide (4NQO) induced head and neck carcinogenesis study

Adult *Iqgap1*^*+/+*^ and *Iqgap1*^*-/-*^ mice (between 6 and 7 weeks of age) were treated with the carcinogen 4NQO (Sigma, St. Louis, MO) in their drinking water at a concentration of 100 μg/ml (stored at 4˚C as a 5 mg/mL stock until use) for a period of 16 weeks. The mice were then returned to normal drinking water for 5 weeks. At the end of this period, mice were euthanized, overt tumors on the tongue and esophagus were quantified, and the tissues were collected for histological analysis.

### Histological analysis

The tongues and esophagus were harvested and fixed in 4% paraformaldehyde (in PBS) for 24 hours, then switched to 70% ethanol for 24 hours, embedded in paraffin, and thin-sectioned (5μm). Every 10^th^ section of the tissues was stained with H&E and examined by a pathologist for the presence of dysplasia and squamous cell carcinomas.

### Immunofluorescence

Tissue sections were deparaffinized in xylenes and rehydrated by 100%, 95%, 70% and 50% ethanol then water. Antigen unmasking was achieved by heating with 10 mM sodium citrate buffer (pH 6) for 20 minutes. After cooling and washes in TBS, slides were blocked with Superblock blocking buffer (ThermoFisher) for 1 hour at room temperature. Primary antibodies used included anti-IQGAP1 (1:100; ab25950, Abcam), anti-pS6 (1:100, #4858, Cell signaling). Slides were incubated in primary antibody at 4˚C, overnight in a humidified chamber. After washes in TBS, Alexa goat-anti-rabbit 594 (Invitrogen) at 1:500 dilution in block was applied for 1 hour at room temperature, followed by TBS washes. Sections were counterstained with DAPI and mounted in Prolong Diamond Antifade Mountant (Invitrogen). All images were taken with a Zeiss AxioImager M2 microscope using AxioVision software version 4.8.2.

### Image analysis and statistical analysis

One-sided Student t-tests were used to determine the significance of AKT, ERK and S6 activation upon *in vivo* EGF stimulation. The percentage of activated protein was calculated by comparing the ratio of phosphorylated protein signal to total protein signal in both PBS injected (control) and EGF injected (stimulated) mice. Protein activation by EGF was then calculated by determining the fold change of percent-activated protein upon EGF (activated protein in EGF stimulated mice/activated protein in PBS treated mice). Error bars (standard deviation) were calculated by taking average of 4 or 6 replicates.

To determine the significance of differences in multiplicity of cancer foci and disease severity in the animal tongue and esophagus, a two-sided Wilcoxon rank sum test was performed for statistical significance using MSTAT statistical software version 6.3.1(http://www.mcardle.wisc.edu/mstat, last accessed January 23, 2018).For disease severity, each cancer grade was represented by a number (well-differentiated cancer=5, moderated-differentiated=6, poorly-differentiated=7) to allow statistical comparison using Wilcoxon rank sum test.

To quantify pS6 levels between groups of mice, 10 microscopic images (20X) of regions of normal, dysplasia and cancer were captured per mouse. All images were taken under constant exposure time. Images were processed using ImageJ software version 10.2 (NIH, Bethesda, MD). pS6 levels on each image were calculated by integrated intensity of pS6 signal, normalized by the DAPI signal from the same area. For statistical analysis, pS6 quantification of the 3 disease stages (normal, dysplasia and cancer) were calculated separately by taking the average of pS6 level in the corresponding 10 images per mouse, and the data was pooled across replicates. A two-sided Student ttest was used to determine the significance of differences in pS6 signal between *Iqgap1*^*+/+*^ and *Iqgap1*^*-/-*^ for each disease stage.

## RESULTS

### Loss of IQGAP1 reduces PI3K signaling and cancer cell survival *in vitro*

To test whether IQGAP1 is required for PI3K signaling in cancer, Iqgap1-knockout cells were generated using CRISPR-Cas9 in the MDA-MB-231 breast cancer cell line. Immunoblotting showed that when IQGAP1 was knocked-out, the levels of activated AKT (represented as pAKT/AKT) was significantly downregulated (Figure 1A), indicating that IQGAP1 is necessary for PI3K signaling in this cancer cell line, which is consistent with results from our previous study [11]. We then tested whether IQGAP1-mediated PI3K signaling is required for growth/survival of HNSCC cells. IQ3 and WW peptide were used to specifically disrupt IQGAP1-mediated PI3K signaling [11]. The addition of either IQ3 or WW greatly inhibited cell growth/survival in both UM-SCC1 and UPCISCC-90 cells, derived from HPV-negative and HPV-positive human HNSCCs, respectively (Figure 1B). These results indicate that IQGAP1 contributes to PI3K signaling pathway in cancer cells, and that IQGAP1-mediated PI3K signaling is necessary for HNSCC cell growth/survival.

**Figure 1.**
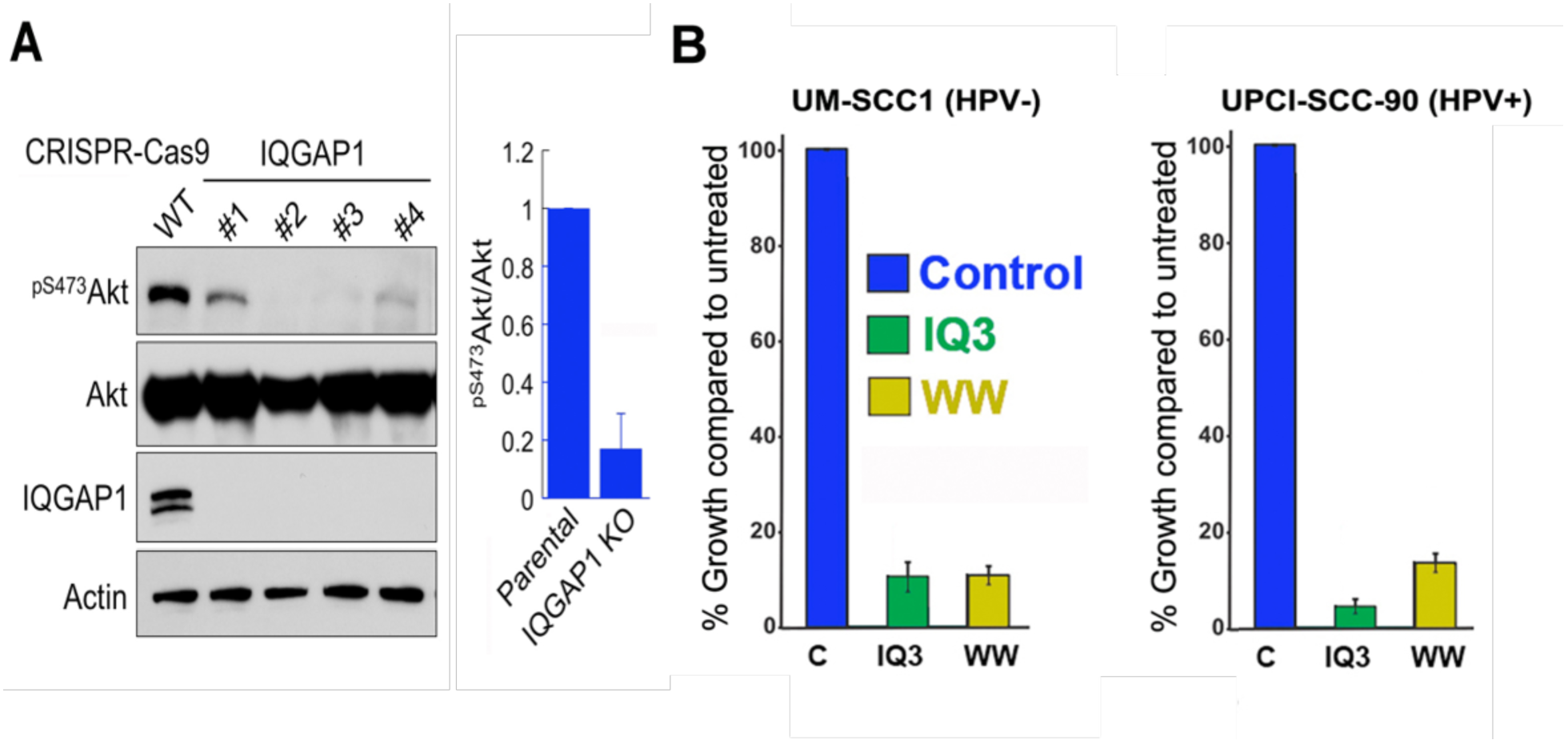
Requirement for IQGAP1 *in vitro*. A) IQGAP1 CRISPR-Cas9 knockout cells were harvested and cell lysates were analyzed by immunoblotting with the indicated antibodies (left). Intensity of immunoblots were quantified with ImageJ and the graph is shown as mean ± SD of the 4 independent IQGAP1-null clones or parental cells (right). B) HNC cells were treated with the indicated peptides at 30μM concentration for 72 hours. The number of viable cells was counted. The graph shows results of three independent experiments.

### IQGAP1 is necessary for efficient PI3K signaling upon EGF-stimulation *in vivo*

Previous studies suggest that IQGAP1 scaffolds both the Ras-MAPK and PI3K signaling pathways [11, 12, 25], which are both implicated in various epithelial cancers. To assess whether IQGAP1 is necessary for activation of PI3K/AKT pathway and Ras-MAPK pathway signaling *in vivo*, we utilized a genetic approach making use of the *Iqgap1* knockout mice (*Iqgap1*^*-/-*^) [21]. Both the PI3K/AKT and Ras-MAPK pathways are activated upon EGF stimulation *in vivo* [23]. EGF (5ug/g weight) was subcutaneously injected into 5-week-old *Iqgap1*^*+/+*^ and *Iqgap1*^*-/-*^ mice and dorsal skin was harvested 10 minutes post-EGF stimulation and subjected to protein analysis by immunoblotting. The activation of PI3K/AKT pathway was scored as the ratio of pAKT/AKT and pS6/S6, while the activation of Ras-MAPK pathway was scored by monitoring the pERK/ERK ratio. We found that AKT activation upon EGF stimulation was significantly lower in *Iqgap1*^*-/-*^ mice compared to *Iqgap1*^*+/+*^ mice (P=0.006, Student t-test, one-sided) (Figure 2A-B). We also observed a significantly lower S6 activation in *Iqgap1*^*-/-*^ mice compared to *Iqgap1*^*+/+*^ mice (P=0.038) (Figure 2A, 2C). The decreased activation of both AKT and S6 in *Iqgap1*^*-/-*^ mice indicated that IQGAP1 is required for efficient activation of PI3K signaling upon EGF stimulation. ERK activation in response to EGF followed a decreasing trend in *Iqgap1*^*-/-*^ mice, but it did not reach the 95% confidence interval (P=0.07, Figure 2A,2D). Interestingly, ERK activation displayed a dramatic sex difference in response to EGF: the absence of IQGAP1 caused a large reduction in ERK activation in female mice, while not affecting ERK activation in male mice (Supplementary Figure 1). These results indicate that IQGAP1 contributes quantitatively to the activation of the PI3K/AKT pathway in response to EGF, and potentially also to Ras-MAPK pathway, albeit to a reduced extent.

**Figure 2.**
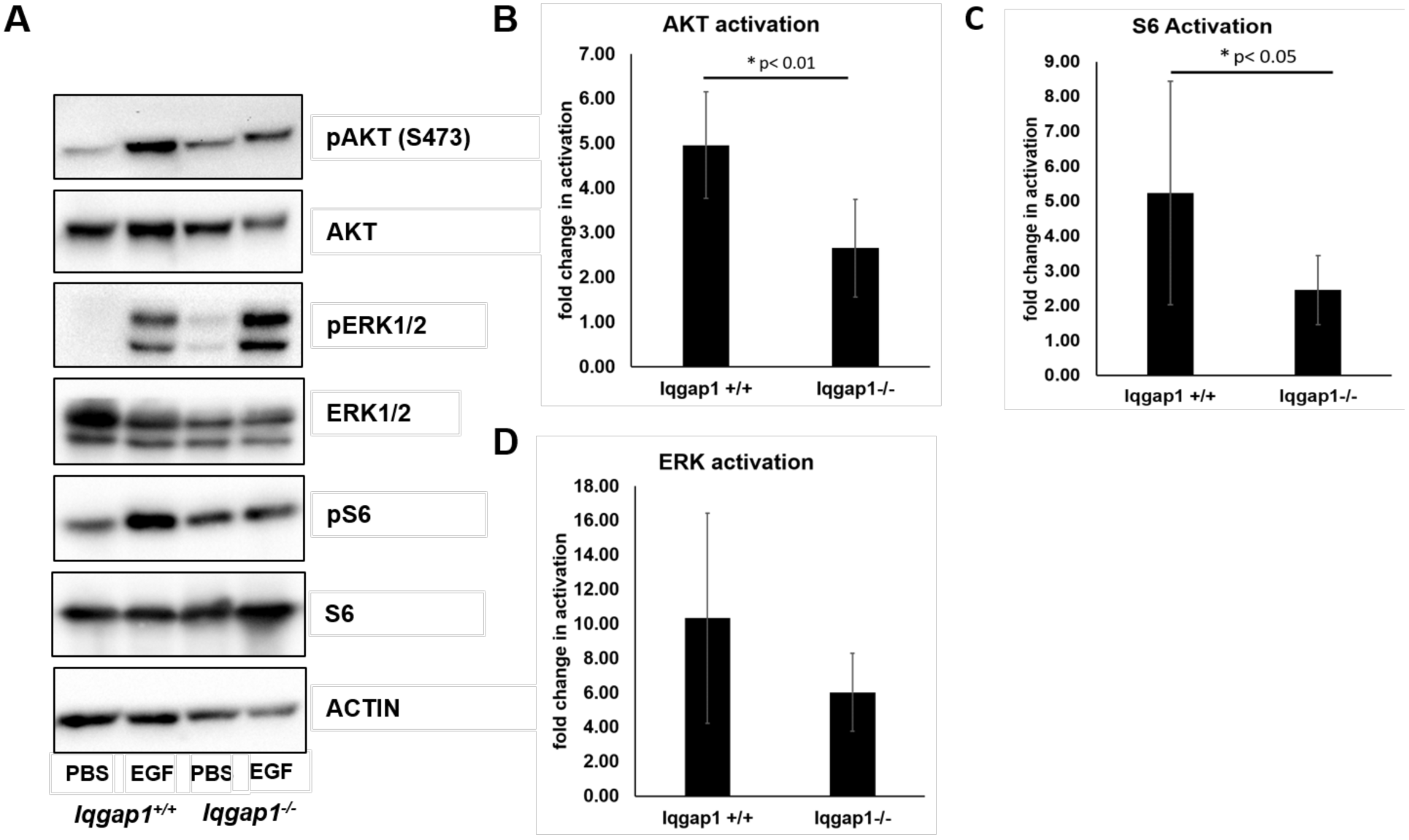
IQGAP1 is necessary for efficient PI3K signaling upon EGF-stimulation *in vivo*. 5-week-old *Iqgap1*^*+/+*^ and *Iqgap1*^*-/-*^ mice were stimulated with subcutaneous injection of EGF (5μg/g weight). PBS only was injected in vehicle control groups. Dorsal skin was harvested 10 minutes later, from which tissue lysates were made and analyzed by immunoblotting. A) Immunoblots detecting pAKT(S473) and total AKT, pERK1/2 and total ERK1/2, p-S6 and total S6 and actin (loading control). The volume of each band (in Fig. 2A) was measured and activation levels of AKT, ERK1/2 and S6 were determined by taking ratios of the (phosphorylated-protein/ total protein) upon EGF stimulation compared to PBS control. The quantified activation levels of (B) AKT, (C) S6 and (D) ERK are shown. Error bars were calculated by calculating the standard deviation of the average of 5 or 6 replicates. T-test was used for statistical analysis.

### Loss of IQGAP1 reduces frequency of high grade head and neck cancers

As shown above (Figs. 1 and 2), and also seen by other’s [11], IQGAP1 is necessary both in cell culture and *in vivo* for efficient PI3K/AKT signaling. The PI3K/AKT pathway is often activated in HNSCC [3], and disruption of IQGAP1-mediated PI3K signaling by the IQ3 peptide decreased HNSCC cell survival (Figure 1B). Therefore, we tested whether loss of IQGAP1 prevents or reduces the onset and/or severity of HNSCC using a well-established mouse model for HNSCC, in which the synthetic oral carcinogen, 4-nitroquinoline 1-oxide (4NQO), is used to drive tumorigenesis [20]. Groups of *Iqgap1*^*+/+*^ (n=19) and *Iqgap1*^*-/-*^ (n=16) mice were given 4NQO (100μg/mL) in their drinking water for 16 weeks, followed by 8 weeks of normal drinking water (Figure 3A). Mouse weight was used as a criterion to monitor mouse health. Morbidity issues, as reflected by weight loss, arose in a few *Iqgap1*^*+/+*^ mice by week 16, the end-point of 4NQO treatment, and worsened as the experiment proceeded. Interestingly, no weight loss was evident in the *Iqgap1*^*-/-*^ group of mice. For humane reasons, we ended the study early, at 21 weeks instead of 24 weeks, because more than 20% of *Iqgap1*^*+/+*^ mice showed significant weight loss.

**Figure 3.**
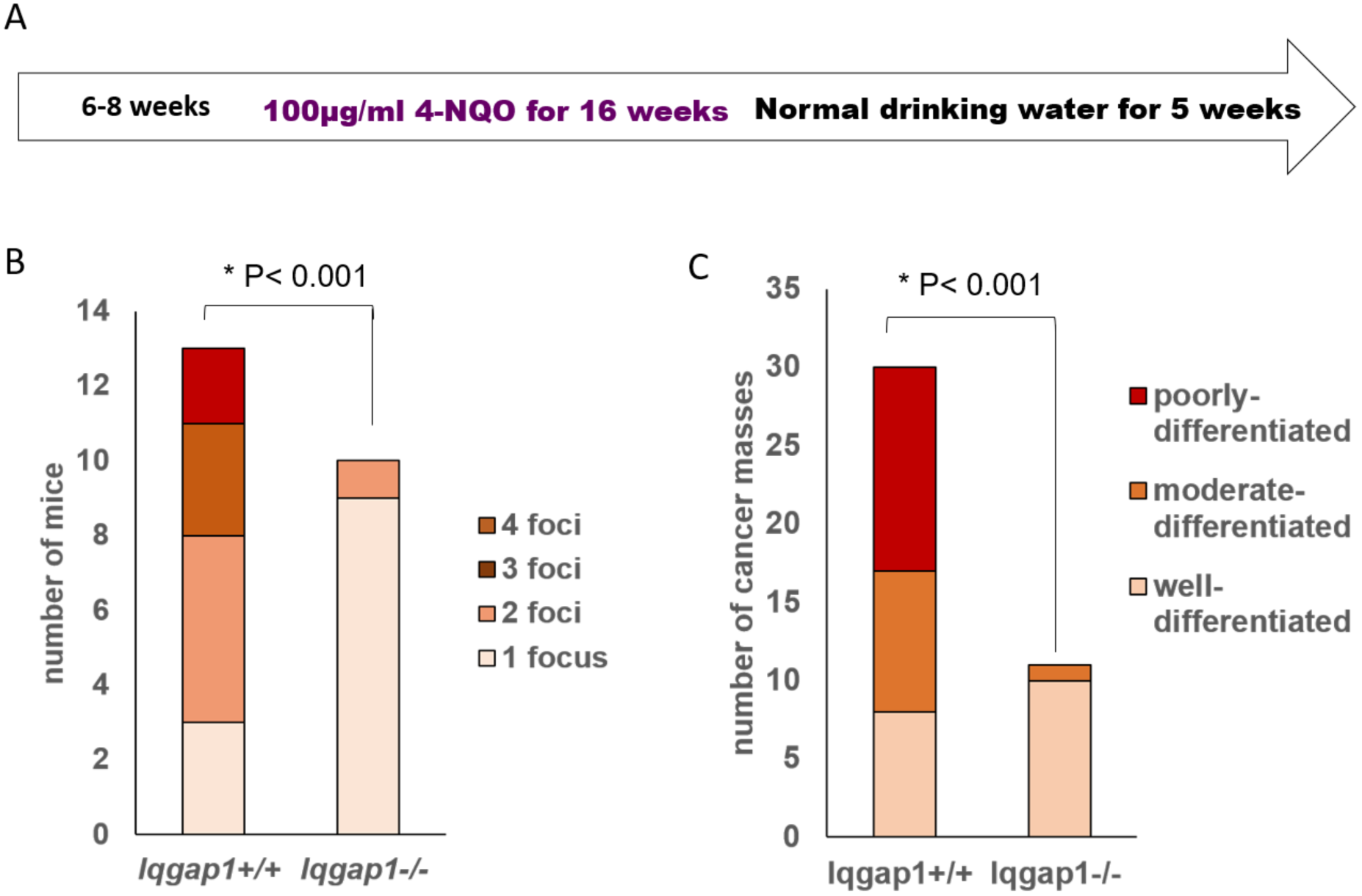
Loss of IQGAP1 significantly reduces the frequency of high-grade head and neck cancers. A) 4NQO treatment timeline. B) Numbers of cancer foci per mouse (combined foci numbers in both esophagus and tongue). C) Numbers of cancer foci with the indicated cancer grade (combined foci number in both esophagus and tongue). Asterisk indicates statistically significance.

At 21 weeks, both *Iqgap1*^*+/+*^ and *Iqgap1*^*-/-*^ groups of mice had 100% incidence of overt tumors in their tongue and esophagus. Disease severity for both tongue and esophagus was assessed by histopathological analysis and cancer grade was assigned. We found that 68.4% of *Iqgap1*^*+/+*^ mice and 62.5% of *Iqgap1*^*-/-*^ mice developed invasive carcinoma, indicating that IQGAP1 did not significantly affect overall percentage of mice with cancers(Table 1, P=0.73, Fisher exact test). However, when we assessed the multiplicity of cancer foci, we found that *Iqgap1*^*+/+*^ mice developed significantly higher numbers of cancer foci per mouse when compared to *Iqgap1*^*-/-*^ mice (Figure 3B, mean value of 2.3 vs. 1.09, p<0.001, Wilcoxon Rank Sum test). We also found that *Iqgap1*^*+/+*^ mice developed significantly higher numbers of poorly-differentiated, high-grade cancer compared to the *Iqgap1*^*-/-*^ mice, which developed mostly well-differentiated, low-grade cancer foci (Figure 3C, mean value of 6.13 vs. 5.09, p<0.001, Wilcoxon Rank Sum test). Altogether, we found there to be a significant reduction in both cancer foci multiplicity and cancer severity in *Iqgap1*^*-/-*^ treated with 4NQO. This demonstrates that IQGAP1 contributes quantitatively to head and neck carcinogenesis.

**Table 1.** **Disease severity in 4NQO-treated *Iqgap1*^*+/+*^ and *Iqgap1*^*-/-*^ mice.**

|  | number of mice | no disease | dysplasia |  |  | carcinoma |  |  |
| --- | --- | --- | --- | --- | --- | --- | --- | --- |
|  |  |  | mild | moderate | severe | well-differentiated | moderate-differentiated | poorly-differentiated |
| *lqgap1*<sup>+/+</sup> | 19 | 1 | 1 | 1 | 4 | 3 | 5 | 4 |
| *lqgap1*<sup>-/-</sup> | 16 | 0 | 1 | 1 | 9 | 4 | 1 | 0 |

**Table 1. Disease severity in 4NQO-treated *lqgap1*<sup>+/+</sup> and *lqgap1*<sup>-/-</sup> mice.**
|  | number of mice | no disease | dysplasia |  |  | carcinoma |  |  |
| --- | --- | --- | --- | --- | --- | --- | --- | --- |
|  |  |  | mild | moderate | severe | well-differentiated | moderate-differentiated | poorly-differentiated |
| *lqgap1*<sup>+/+</sup> | 19 | 0 | 3 | 2 | 4 | 5 | 2 | 3 |
| *lqgap1*<sup>-/-</sup> | 16 | 0 | 3 | 1 | 7 | 5 | 0 | 0 |

### IQGAP1 protein levels are upregulated in HNSCC

IQGAP1 is overexpressed in many human cancers [15, 18, 26, 27]. We therefore looked at levels of IQGAP1 protein in our 4NQO-induced HNSCC model. Based upon immunofluorescence, higher levels of IQGAP1 protein were detected in cancer regions compared to normal regions of both the tongue and the esophagus of *Iqgap1*^*+/+*^ mice (Figure 4). Tissue from *Iqgap1*^*-/-*^ mice, served as negative controls (Figure 4). The upregulation of IQGAP1 in 4NQO-induced HNSCC in mice is therefore consistent with the previous study reporting overexpression of IQGAP1 in HNSCC patients [18].

**Figure 4.**
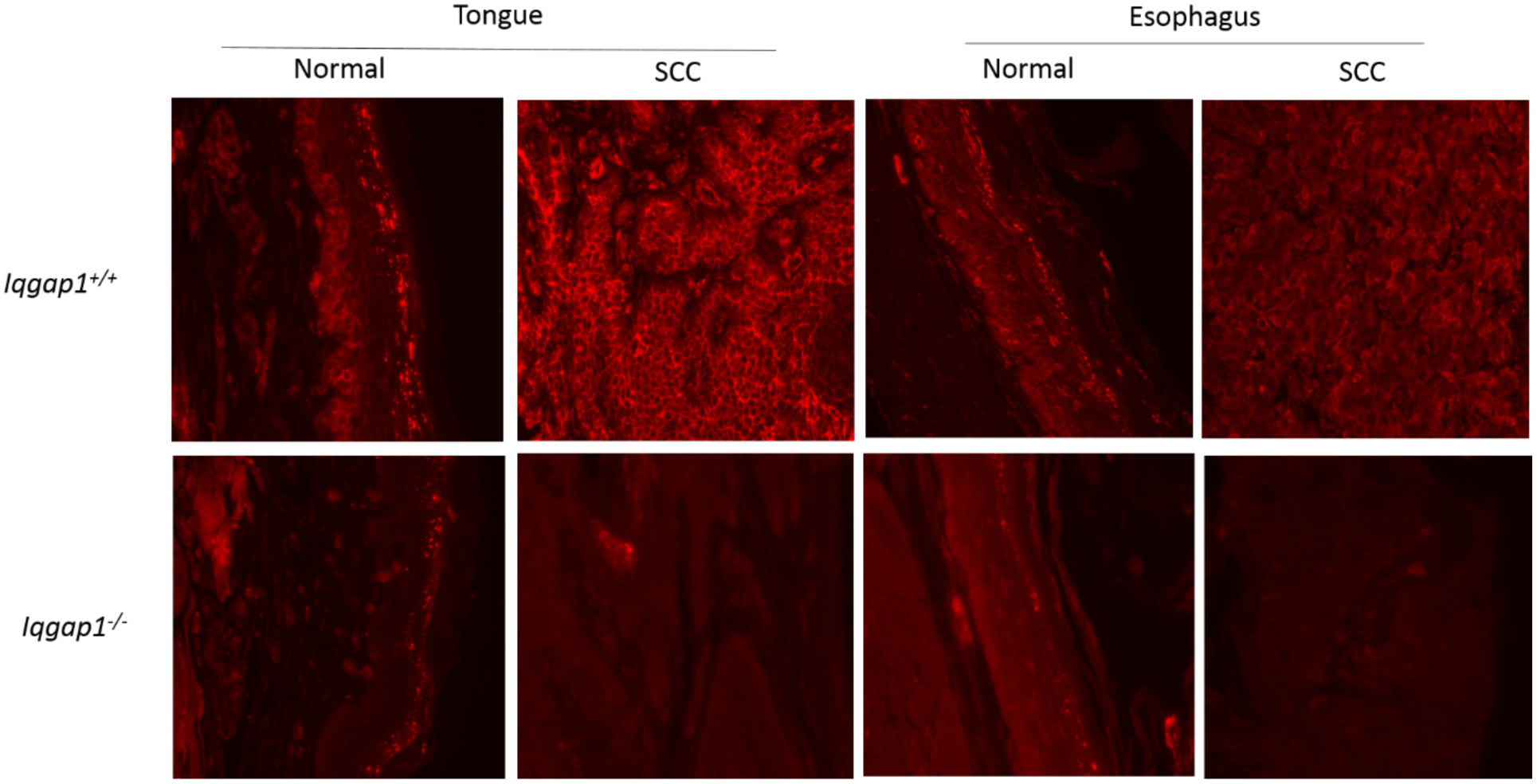
IQGAP1 protein levels are upregulated in SCC in both tongue and esophagus tissue of *Iqgap1*^*+/+*^ mice. Higher levels of IQGAP1 (red, located on cell membrane; levels of protein indicated by intensity of the signal) were detected in tongue cancer as well as esophagus cancer in *Iqgap1*^*+/+*^ mice, when compared to the normal regions. As expected, no cell membrane signals of IQGAP1 were detected in *Iqgap1*^*-/-*^ mice. All images were taken under consistent exposure time.

### PI3K signaling is downregulated in IQGAP1-null mice as disease progresses upon 4NQO treatment

IQGAP1 contains multiple protein-interacting domains that allow IQGAP1 to regulate diverse cellular pathways, such as scaffolding and facilitating Ras-MAPK pathway and PI3K-AKT pathway, promoting Wnt signaling by enhancing nuclear localization of βcatenin, among others [11-13, 17]. We wanted to further dissect which of these pathways contributing to IQGAP1-mediated head and neck carcinogenesis. Given that 4NQO is an EGFR-driven cancer model [20], in addition to previous data that supports the loss of efficient PI3K signaling with the absence of IQGAP1 *in vivo* and *in vitro*, we first looked at whether PI3K signaling was downregulated in *Iqgap1*^*-/-*^ experimental mice during disease progression. In the esophagus, pS6 levels remained consistent during disease progression in *Iqgap1*^*+/+*^ mice, but decreased in *Iqgap1*^*-/-*^ mice (Figure 5). The 4NQO-induced cancers arising in the *Iqgap1*^*-/-*^ mice had a significantly lower level of pS6 than the cancers arising in the *Iqgap1*^*+/+*^ mice (Figure 5, P=0.02, Student t test, two-sided). The reduced pS6 in esophageal cancer lacking IQGAP1 are consistent with the hypothesis that IQGAP1 is contributing to 4NQO-induced HNSCC by driving PI3K signaling. However, IQGAP1 status did not correlate significantly with pS6 levels in tongue cancer (Supplementary Figure 2), indicating that IQGAP1 may exhibit tissue-specific functions in its ability to promote carcinogenesis.

**Figure 5.**
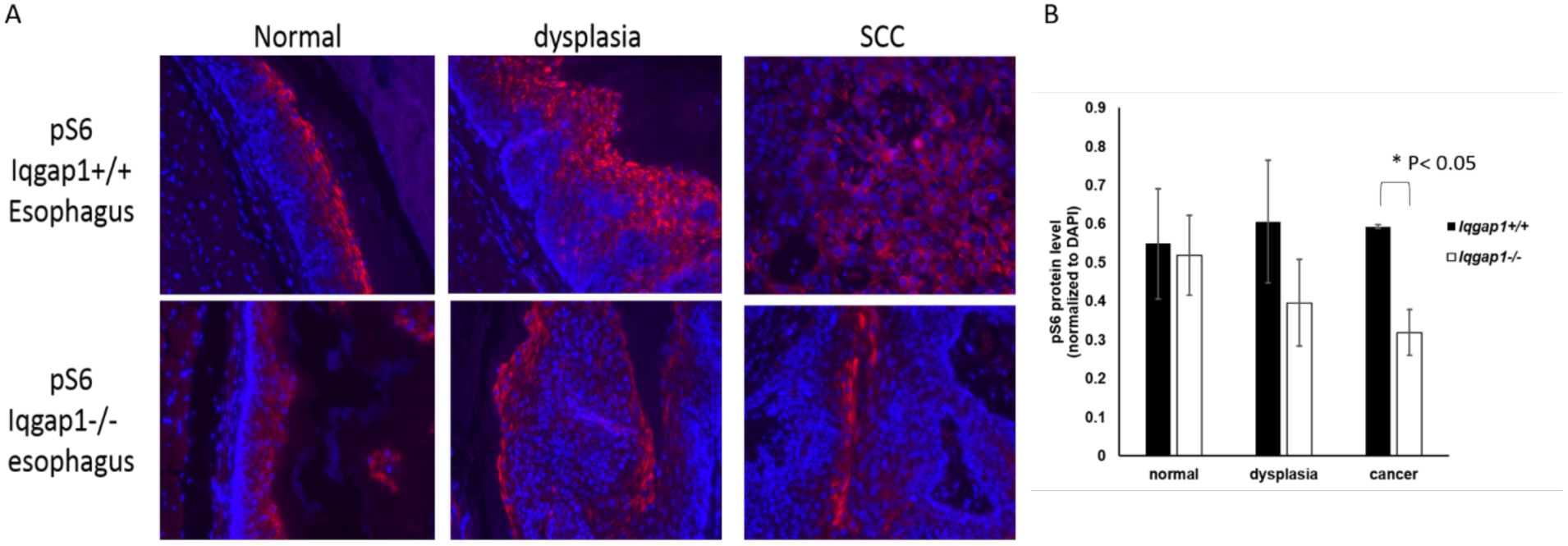
Loss of IQGAP1 reduces PI3K signaling in 4NQO-induced HNSCC. A) Immunofluorescence on 4NQO-treated esophagus (normal, dysplasia, cancer). Red: pS6, blue: DAPI. B) Quantification of pS6 levels in different disease stages. Asterisk indicates statistically significance (P<0.05).

## DISCUSSION

In this report, we demonstrate that IQGAP1 is necessary for growth/survival of human head and neck cancer cells *in vitro*, and contributes quantitatively to the incidence and severity of HNSCC disease in an *in vivo* mouse model for head and neck carcinogenesis. Specifically, cells knocked out for IQGAP1 showed reduced PI3K signaling *in vitro* (Figure 1A). Likewise, IQGAP1 was found to contribute quantitively to PI3K signaling in response to EGF stimulation *in vivo* in *Iqgap1*^*-/-*^ mice (Figure 2). Disruption of IQGAP1-mediated PI3K signaling by IQ3 inhibited head and neck cancer cell survival *in vitro* (Figure 1B). And *in vivo*, utilizing the 4NQO-induced head and neck cancer model known to be dependent upon EGFR signaling [20], *Iqgap1*^*-/-*^ mice showed reduced frequency of high grade cancer and reduced number of cancer foci (Figure 3), indicating for the first time that IQGAP1 contributes to HNSCC. IQGAP1 protein level were upregulated in cancer foci in both tongue and esophagus of 4NQO-treated *Iqgap1*^*+/+*^ mice (Figure 4), consistent with what is seen in human head and neck cancer [18] and PI3K signaling in the 4NQO-induced esophageal cancers was downregulated in the *Iqgap1*^*-/-*^ mice compared to the *Iqgap1*^*+/+*^ mice (Figure 5), consistent with the hypothesis that IQGAP1 is acting through PI3K signaling to promote HNSCC.

Upon 4NQO treatment, we saw significant differences in the multiplicity and disease severity of cancer foci, in which the *Iqgap1*^*-/-*^ experimental group developed fewer cancer foci per mouse as well as fewer of high-grade cancers. However, we did not observe a significant difference in overt tumor incidence (100% in both groups) and percentage of mice with cancer (68.4% of *Iqgap1*^*+/+*^ vs. 62.5% of *Iqgap1*^*-/-*^, P=0.73). It is possible that the high dose of 4NQO used in this study masked some effects of IQGAP1.

Our observation of early rising morbidities, particularly in the *Iqgap1*^*+/+*^ mice support the notion that the dose of carcinogen could be masking effects of IQGAP1 status on the percentage of mice that develop overt tumors and with cancers. The genetic background of our experimental mice could also contribute to the masking effects of the high dosage of 4NQO. Previous research using same 4NQO treatment followed by 12 weeks of normal drinking water, caused 100% incidence of invasive squamous cell carcinoma in *C57Bl/6* mice [20]. The experimental mice used in our study were on a mixed genetic background of 50%*FVB*/50%(*129+C57Bl/6*). *FVB* mice have been shown to be highly susceptible to chemical carcinogen-induced squamous cell carcinoma in the skin compared to other strains of mice [28]. Therefore, it is possible that the 4NQO treatment condition was too harsh for this mixed genetic background, as evidenced by morbidity issues, and that the response to the 4NQO treatment masked the extent of the effect of loss of IQGAP1 on tumorigenicity.

We showed that upon 4NQO treatment, esophageal PI3K signaling in *Iqgap1*^*-/-*^ mice was downregulated as the disease progressed, and significantly lower in cancer when compared to *Iqgap1*^*+/+*^ mice. This supports the hypothesis that IQGAP1 promotes HNSCC by driving PI3K signaling, given that IQGAP1 scaffolds PI3K signaling [11]. However, no significant difference was observed for PI3K signaling in tongue (Supplemental Figure 2), suggesting that there may be site-specific differences in how IQGAP1 contributes to HNSCC. It is also worth noting that more tumors developed in the esophagus than in the tongue. A possible hypothesis for this observation is that, in the tongue of Iqgap1^-/-^ mice, another factor acts to maintain signaling when IQGAP1 is absent. One candidate for this other factor is IQGAP3, another member of the IQGAP family that has highly homologous structural motifs to IQGAP1 [29, 30]. IQGAP3 functions similarly to IQGAP1, including its interactions with Rac1 and Cdc42 for actin filament regulation, and promoting Ras/ERK signaling [24, 30, 31]. Interestingly, when IQGAP1 was knocked down by siRNA in human hepatocellular cancer cells, levels of IQGAP3 protein were upregulated [32]. Although it is still unknown whether IQGAP3 scaffolds for PI3K signaling, considering its similarity to IQGAP1, it is possible that under certain circumstances, IQGAP3 compensates and maintains PI3K signaling when IQGAP1 is absent. In our mice skin lysates, we did not observe a significant change on IQGAP3 levels in tissues null for IQGAP1 (Supplementary figure 1A, E). But this does not exclude the possibility that such compensation happens in tongue epithelia in mice.

Although we have focused on IQGAP1-mediated PI3K signaling, we cannot discount the possibility that IQGAP1 may act through other pathways to promote HNSCC. *Jameson et. al.* proposed that, under DMBA-TPA treatment - a Ras-driven skin cancer model, *Iqgap1*^*-/-*^ mice were resistant to skin tumorigenesis, and this reduced tumorigenesis was due to the lack of IQGAP1-scaffolded Ras-MAPK signaling in the absence of IQGAP1 [12]. In our study, we utilized the 4NQO-induced HNSCC model, which is an epidermal growth factor receptor (EGFR)-driven tumorigenesis model [20]. Considering that both PI3K and Ras-MAPK signaling are downstream of EGFR, it remains possible that IQGAP1 acts through Ras-MAPK pathway when promoting HNSCC. Multiple studies have linked IQGAP1 with Wnt*/* β-catenin signaling in cancer promotion [33-35]. IQGAP1 is required for sufficient R-spondin (RSPO)-induced Wnt*/* βcatenin signaling [33]. In thyroid cancer, knockdown of IQGAP1 reduced Wnt/β-catenin signaling both *in vitro* and in xenograft model, inhibiting tumor growth and epithelial-mesenchymal transition (EMT) marker expression [34]. In hepatocellular carcinoma cell line, IQGAP1 co-immunoprecipitated with β-catenin, increased β-catenin expression and nuclear β-catenin level, and activated downstream transcription [35]. Future studies such as RNAseq or Mass-Spectrometry based proteomic analysis will shed more light on whether other IQGAP1-mediated pathways may be involved in promoting HNSCC.

## ACKNOWLEDGMENTS

This work was supported by funds from NIH to P.F.L. (R35-CA210807, P01-CA022443), R.A.A. (R01-GM57549, R01-CA104708) and A.C.R. (R01-CA163662), funds to P.F.L and A.C.R. from the UW Head and Neck Cancer SPORE (P50-DE026787), and funds to P.F. L., A.C.R. and R.A.A. from the UW School of Medicine and Public Health and the UW Carbone Cancer Center (UWCCC). This study made use of UWCCC shared services, which are supported by an NCI Cancer Center grant (P30-CA014520). We acknowledge David Sacks (NIH) for the *Iqgap1*-null mice.

## Authorship Contributions

T.W. participated in concept design, performed experiments and wrote the manuscript; S.C. performed experiments and contributed to the writing the manuscript; D.B. performed pathological analyses; A.C.R and R.A.A participated in concept design and critique of the manuscript, and P.F.L. oversaw concept design and participated in writing the manuscript.

